# LaeA controls citric acid production through regulation of a citrate exporter encoding *cexA* in *Aspergillus luchuensis* mut. *kawachii*

**DOI:** 10.1101/748426

**Authors:** Chihiro Kadooka, Eri Nakamura, Kazuki Mori, Kayu Okutsu, Yumiko Yoshizaki, Kazunori Takamine, Masatoshi Goto, Hisanori Tamaki, Taiki Futagami

## Abstract

The putative methyltransferase LaeA is a global regulator of metabolic and development processes in filamentous fungi. We characterized the homologous *laeA* genes of the white koji fungus *Aspergillus luchuensis* mut. *kawachii* (*A. kawachii*) to determine their role in the hyperproduction of citric acid. The Δ*laeA* strain exhibited a significant reduction in citric acid productivity. Cap-analysis gene expression (CAGE) revealed that *laeA* is required for the gene expression of a putative citrate exporter encoding *cexA*, which is critical for citric acid production. The deficit in the productivity of citric acid by the Δ*laeA* strain was rescued by the overexpression of *cexA* to a level comparable with that of the *cexA*-overexpressing Δ*cexA* strain. In addition, chromatin immunoprecipitation coupled with quantitative PCR (ChIP-qPCR) analysis indicated that LaeA regulates the expression of *cexA* via methylation levels of the histones H3K4 and H3K9. These results indicate that LaeA is involved in citric acid production through epigenetic regulation of *cexA* in *A. kawachii*.

**IMPORTANCE:** *A. kawachii* has been traditionally used for production of the distilled spirit shochu in Japan. Citric acid produced by *A. kawachii* plays an important role in preventing microbial contamination during the shochu fermentation process. This study characterized homologous *laeA* genes; using CAGE, complementation test, and ChIP-qPCR, it was found that *laeA* is required for citric acid production though regulation of *cexA* in *A. kawachii*. The epigenetic regulation of citrate production elucidated in this study will be useful for controlling the fermentation processes of shochu.

---

Shochu is a traditional Japanese distilled spirit (1). The black koji fungus *Aspergillus luchuensis* and its albino mutant white koji fungus *Aspergillus luchuensis* mut. *kawachii* (*A. kawachii*) are primarily used for the production of shochu. *A. luchuensis* and *A. kawachii* produce enzymes that decompose the starch contained in ingredients such as rice, barley, buckwheat, and sweet potato (2). In addition, they also produce a large amount of citric acid during the fermentation process to prevent the microbial contamination.

*A. luchuensis* is phylogenetically related to *A. niger*, which has been used for industrial citric acid fermentation (3–5). Studies investigating the production of citric acid have been performed in *A. niger* with respect to various aspects (6–8). In previous studies, non-acidifying mutant strains of *A. niger* were analyzed and the *laeA* gene was found to play a significant role in the production of citric acid and secondary metabolites (9). In addition, *laeA* disruption also caused a significant decrease in the citric acid production by *Aspergillus carbonarius*, a species closely related to *A. niger* (10). LaeA was initially identified as a regulator of secondary metabolism in *Aspergillus* spp. (11). Subsequently, LaeA has been primarily studied as a regulator of secondary metabolic and development processes in filamentous fungi (12, 13). A transcriptomic study also supported that *laeA* overexpression and disruption caused the production of secondary metabolites to dramatically change in *A. niger* (14). However, why LaeA is required for citric acid production in *A. niger* and *A. carbonarius* remains to be determined.

In this study, we characterized three *laeA* homologous genes, namely, *laeA*, *laeA2*, and *laeA3*, to determine the regulatory mechanism underlying citric acid production in *A. kawachii*. Study of gene disruption indicated that *laeA* significantly reduced citric acid production; therefore, we further analyzed LaeA-dependent gene expression by cap-analysis gene expression (CAGE) and found that *laeA* is required for expression of a putative citrate exporter encoding *cexA*, which plays a crucial role in citric acid production (15, 16). Further analysis via complementation test and chromatin immunoprecipitation coupled with quantitative PCR (ChIP-qPCR) indicated that LaeA is required for citric acid production through epigenetic regulation of *cexA* in *A. kawachii*.

## RESULTS AND DISCUSSION

### LaeA-like putative methyltransferases in *A. kawachii*

Protein BLAST analysis (https://www.ncbi.nlm.nih.gov/) using the amino acid sequences of *A. nidulans* LaeA (AN0807) and *A. niger* LaeA (An01g12690) as search queries identified three LaeA-like putative methyltransferases in *A. kawachii* using criteria of >50% identity and E-values of <1.00E-100: AKAW_04823, AKAW_11001, and AKAW_06012. AKAW_04823 is extremely similar to *A. niger* LaeA with 94% identity over 375 amino acid residues, likely owing to the close phylogenetic relationship between *A. kawachii* and *A. niger*. In addition, all three *A. kawachii* proteins showed a reciprocal best BLAST hit to *A. nidulans* LaeA, and alignments of these proteins demonstrated that structural homology with the *S*-adenosylmethionine (SAM) binding motif (11) is highly conserved (Fig. 1A). Thus, AKAW_04823, AKAW_11001, and AKAW_06012 as LaeA-like putative methyltransferases were termed LaeA, LaeA2, and LaeA3, respectively.

**Figure 1.**
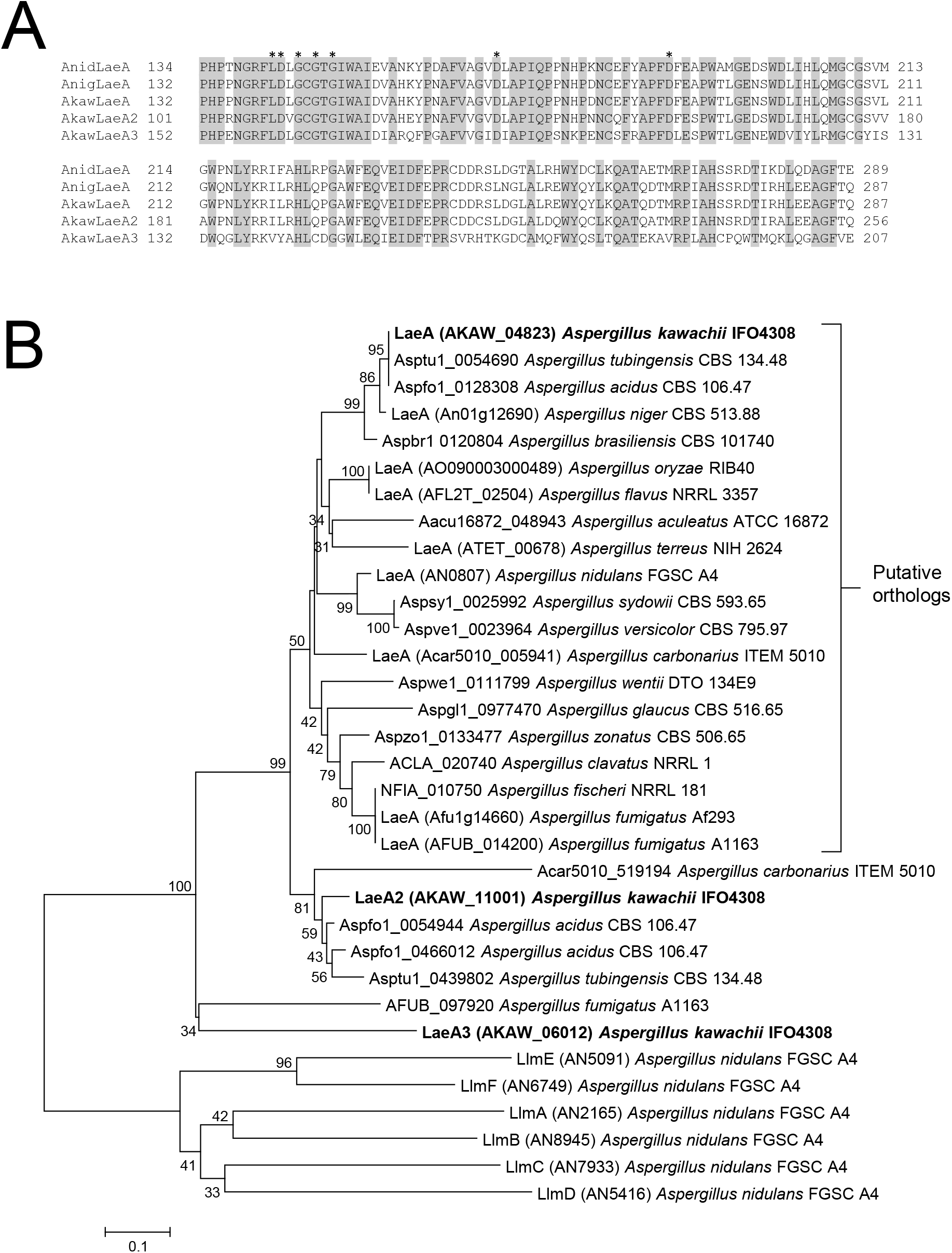
(A) Putative methyltransferase domain sequences of *A. nidulans* (Anid) LaeA, *A. niger* (Anig) LaeA, and predicted *A. kawachii* (Akaw) LaeA-like proteins. The domains were identified by Pfam (https://pfam.xfam.org/). Sequence alignment was performed using ClustalW program in the BioEdit Sequence Alignment Editor (http://www.mbio.ncsu.edu/BioEdit/bioedit.html). Putative intron sequences of *A. kawachii laeA* and *laeA2* genes were confirmed by RNA-seq (data not shown) and translated to the amino acid sequences. Asterisk indicates a putative SAM binding site (11). (B) Phylogenetic tree of amino acid sequences of LaeA-like methyltransferases identified from the *Aspergillus* genome database (http://www.aspgd.org/). The tree was constructed via neighbor-joining method with complete gap deletion on MEGA version 6.0 (36). Bootstrap values (1000 replicates) are indicated at the branches. Low bootstrap values (<25) were removed.

To further clarify the relationship between LaeA, LaeA2, and LaeA3, we performed a phylogenetic analysis of LaeA-like methyltransferases conserved in *Aspergillus* spp. (Fig. 1B). The data set for analysis was obtained by protein BLAST of the *Aspergillus* Genome Database (AspGD; http://www.aspgd.org/) using the amino acid sequences of *A. kawachii* LaeA, LaeA2, and LaeA3 as search queries. *A. kawachii* LaeA is classified into the putative LaeA orthologs group. Conversely, LaeA2 and LaeA3 were assigned to different positions; however, they remain closer to LaeA than to *A. nidulans* Llm (LaeA-like-methyltransferase) proteins (17). This result supports the hypothesis that LaeA2 and LaeA3 are also LaeA-like methyltransferases. LaeA-like methyltransferases, similar to *A. kawachii* LaeA2, were conserved in the *Aspergillus* section *Nigri* (*A. carbonarius* ITEM 5010, *Aspergillus acidus* CBS 106.47, and *Aspergillus tubingensis* CBS 134.48), with the exception of *A. niger* CBS 513.88. In addition, LaeA-like methyltransferases similar to *A. kawachii* LaeA3 were only identified in *A. fumigatus* A1163.

### Colony formation and citric acid production of control, Δ*sC*Δ*laeA*, Δ*sC*Δ*laeA2*, and Δ*sC*Δ*laeA3* strains

To investigate the role of LaeA-like methyltransferases in citric acid production of *A. kawachii*, we constructed each gene disruptant and observed their colony formation (Fig. 2A). For comparison with the respective disruption, here the *A. kawachii* control strain was defined to show the same auxotrophic background.

**Figure 2.**
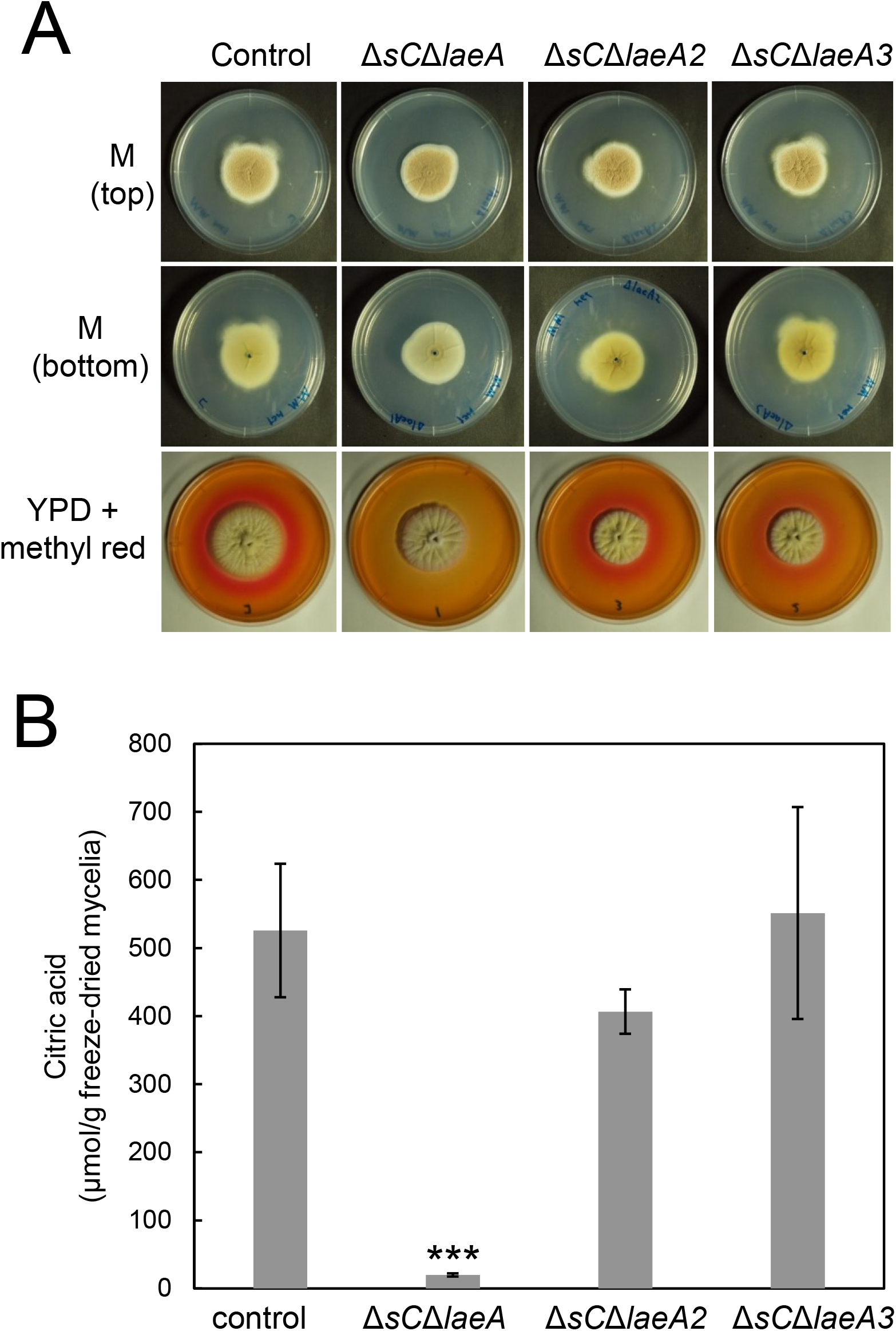
(A) Colony formation of *A. kawachii* control, Δ*sC*Δ*laeA*, Δ*sC*Δ*laeA2*, and Δ*sC*Δ*laeA3* strains. Strains were cultured at 30°C for 5 days on M medium or YPD medium containing methyl red. Agar medium was inoculated with 1 × 10^4^ conidiospores. Methyl red was added as a pH indicator, which turned red at ≤ pH 4.4. (B) Extracellular citric acid production by *A. kawachii* strains; control, Δ*sC*Δ*laeA*, Δ*sC*Δ*laeA2*, and Δ*sC*Δ*laeA3* strains were precultured in M medium for 36 h, then transferred to CAP medium for 48 h. Mean and standard deviation were determined from the results of 3 independent cultivations. Asterisks indicate a statistically significant difference (***, *P* value < 0.001; Student’s t-test) from the results for the control strain.

The Δ*sC*Δ*laeA* strain exhibited colony morphology similar to that of the control strain, but its color was paler than that of control, Δ*sC*Δ*laeA2*, and Δ*sC*Δ*laeA3* strains upon observing the bottom of the minimal (M) medium agar plate (Fig. 2A). This result indicates that LaeA positively regulates the production of the hyphal yellow pigment in *A. kawachii* in addition to reports showing that LaeA is involved in secondary metabolism in *A. niger* (9, 14). However, a difference was observed in the *A. kawachii* Δ*sC*Δ*laeA* strain in that the *A. niger* Δ*laeA* strain showed yellowish mycelia compared with that of the *A. niger* control strain on minimal medium agar plate (9).

In addition, red color surrounding the colonies disappeared only with the Δ*sC*Δ*laeA* strain in the YPD medium containing methyl red as a pH indicator, indicating that the Δ*sC*Δ*laeA* strain shows a non-acidifying phenotype (Fig. 2A). Therefore, we compared the organic acid productivities of control, Δ*sC*Δ*laeA*, Δ*sC*Δ*laeA2*, and Δ*sC*Δ*laeA3* strains to further investigate the role of LaeA-like methyltransferases in citric acid production (Fig. 2B). The strains were precultured in M medium at 30°C for 36 h and then transferred to citric acid production (CAP) medium and further cultured at 30°C for 48 h. Next, citric acid levels in the culture supernatant were measured as the extracellular fraction. CAP medium was used for evaluating organic acid production as it contains a high concentration of carbon source [10% (wt/vol) glucose] as well as appropriate trace elements (6–8).

Compared with the control strain, the Δ*sC*Δ*laeA* strain exhibited a 0.04-fold lower citric acid production, whereas Δ*sC*Δ*laeA2* and Δ*sC*Δ*laeA3* strains exhibited similar level. This result indicates that LaeA plays a significant role in the citric acid production of *A. kawachii*. In addition, this result is consistent with the results of previous reports that LaeA is required for citric acid accumulation in the culture supernatant of *A. niger* and *A. carbonarius* (9, 10).

### Complementation test of Δ*laeA* strain with wild type *laeA*

Because *laeA* disruption caused a significant decline in citric acid production by *A. kawachii*, we examined the complementation of the wild type *laeA* gene in the Δ*sC*Δ*laeA* strain using a *sC* marker. Control, Δ*laeA*, and Δ*laeA*+*laeA* strains were cultured as described above and culture supernatant and mycelia were separated as the extracellular and intracellular fractions, respectively. In the extracellular fraction, the Δ*laeA* strain exhibited 0.08-fold lower citric acid levels than the control strain (Fig. 3A). Conversely, decreases in the production of malic acid and oxoglutaric acid (0.64-fold and 0.70-fold, respectively) were not statistically significant. Decreases in the production of citric acid, malic acid, and oxoglutaric acid were rescued by the complementation of the wild type *laeA* gene.

**Figure 3.**
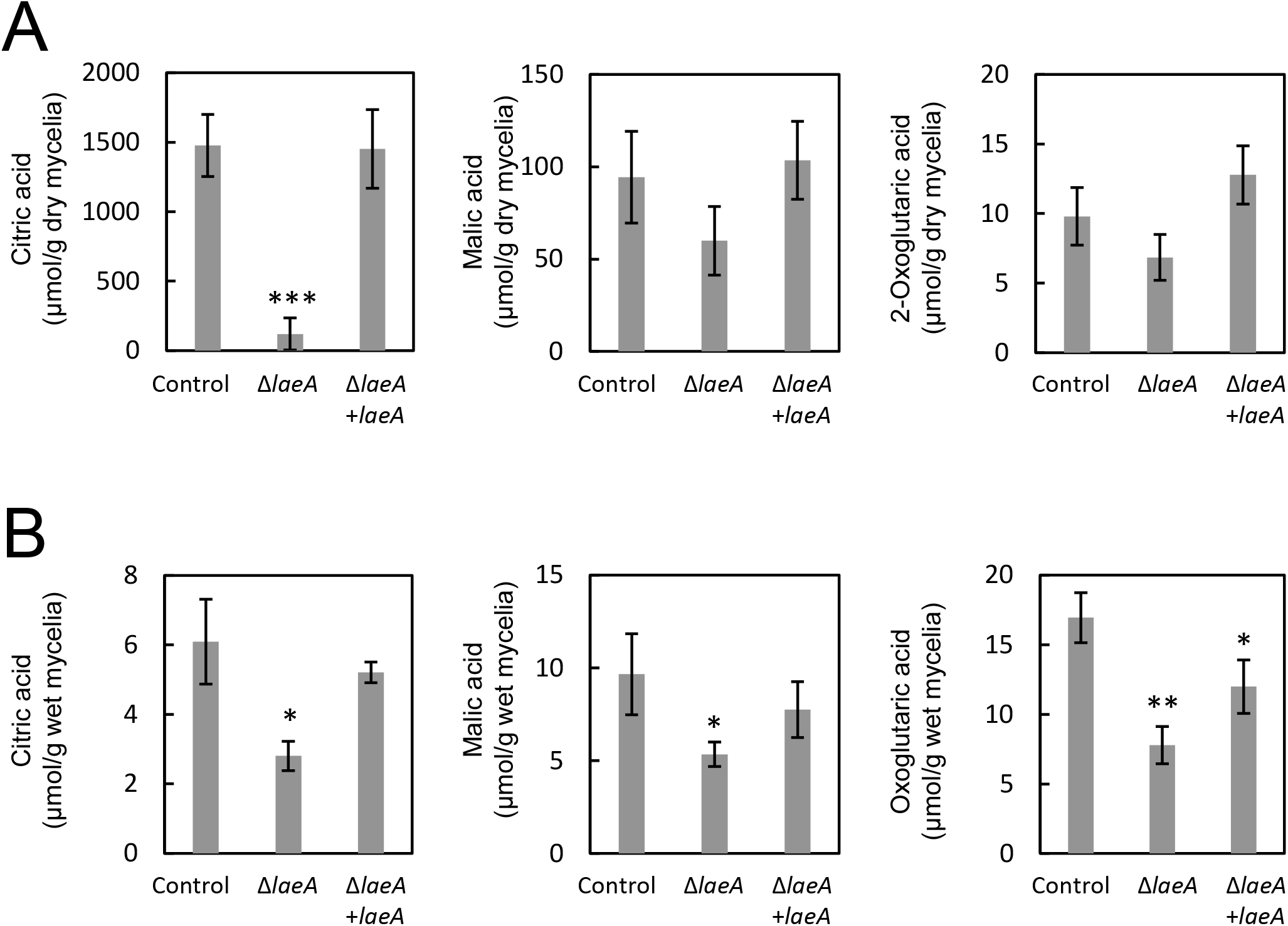
Extracellular (A) and intracellular (B) organic acid production by *A. kawachii* strains. The control, Δ*laeA*, Δ*laeA*+*laeA* strains were precultured in M medium for 36 h, then transferred to CAP medium for 48 h. Mean and standard deviation were determined from the results of 3 independent cultivations. Asterisks indicate statistically significant difference (*, *P* value < 0.05; **, *P* value <0.01; ***, *P* value <0.001; Student’s t-test) from the results for the control strain.

Additionally, in the intracellular fraction, the Δ*laeA* strain exhibited productivities of 0.46-fold lower citric acid, 0.55-fold lower malic acid, and 0.46-fold lower oxoglutaric acid (Fig. 3B). These decreases in organic acid production were rescued by the complementation of the wild type *laeA* gene, although the Δ*laeA+laeA* strain still exhibited 0.71-fold lower oxoglutaric acid production than that of the control strain.

These results indicate that LaeA plays a significant role both in extracellular and intracellular organic acid production in *A. kawachii*. In addition, extracellular citric acid accumulation was most significantly affected by *laeA* disruption.

### Colony formation of control, Δ*laeA*, and Δ*laeA*+*laeA* strains

To explore the physiologic roles of LaeA, we characterized the colony morphology of *A. kawachii* control, Δ*laeA*, and Δ*laeA*+*laeA* strains (Fig. 4A). The Δ*laeA* strain exhibited slightly smaller average colony diameter than those of the control and Δ*laeA*+*laeA* strains on M medium at all tested temperatures at pH 6.5. This growth deficiency was restored at pH 3, whereas it worsened at pH 10.

**Figure 4.**
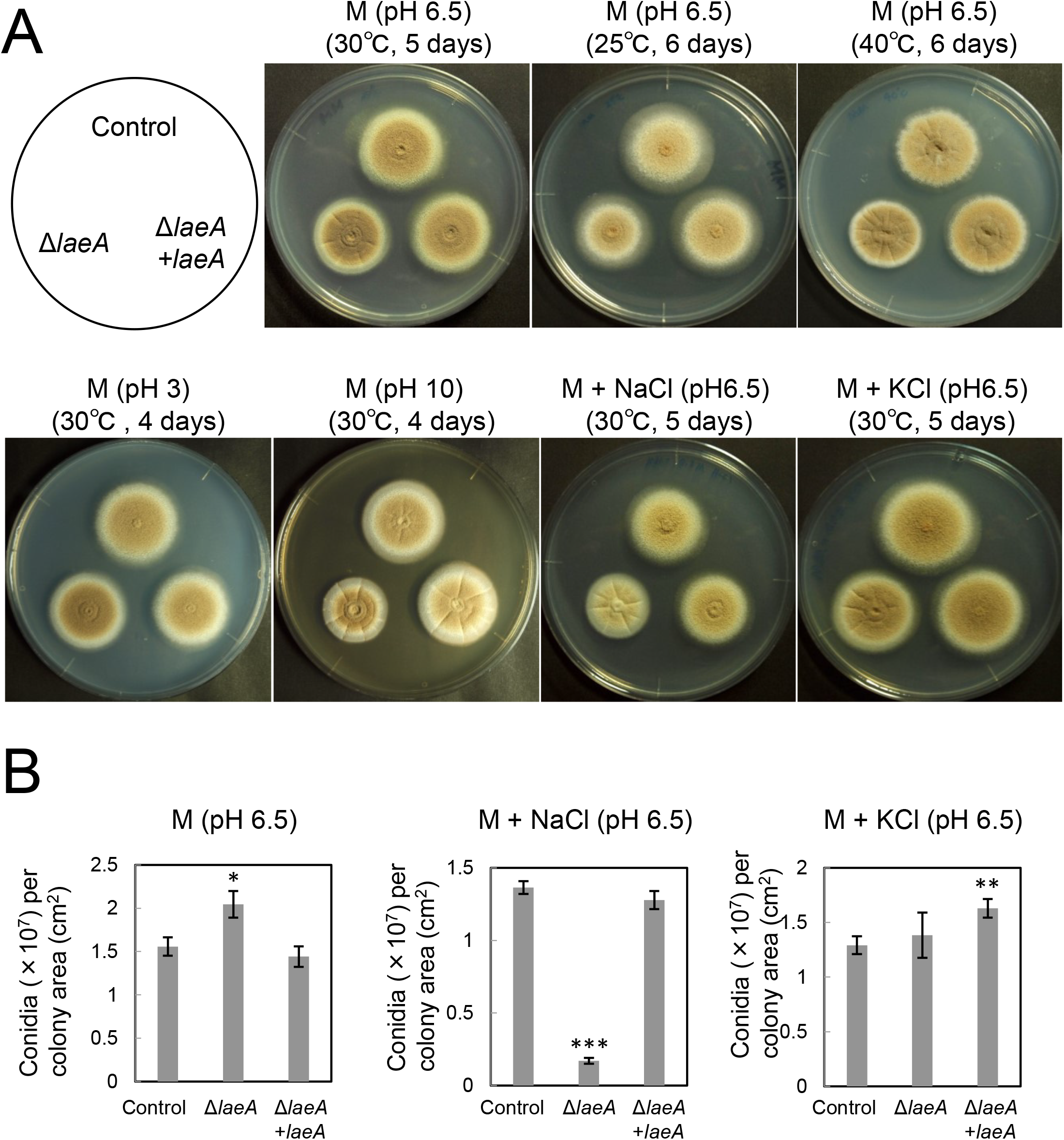
Strains were cultured on M medium with or without stress conditions, and agar medium was inoculated with 1 × 10^4^ conidia. (A) Colony morphology of *A. kawachii* control, Δ*laeA*, Δ*laeA*+*laeA* strains. (B) Conidiation of *A. kawachii* control, Δ*laeA*, Δ*laeA*+*laeA* strains. All results are expressed as mean with standard deviations. Asterisks indicate a statistically significant difference (*, *P* value < 0.05; **, *P* value <0.01; ***, *P* value <0.001; Student’s t-test) from the results for the control strain.

In addition, colonies of the Δ*laeA* strain were paler than those of the control and Δ*laeA*+*laeA* strains in the M medium with 0.8 M sodium chloride but not in that with 0.6 M potassium chloride (Fig. 4A). Thus, we next assessed conidia formation (Fig. 4B). Strains were cultured on M medium with or without 0.8 M sodium chloride or 0.6 M potassium chloride at 30°C for 5 days, at which time the number of conidia formed was determined. The number of conidia per square centimeter of the Δ*laeA* strain was significantly reduced to approximately 0.13-fold of the number formed by the control strain in the M medium containing 0.8 M sodium chloride. Alternatively, the number of conidia per square centimeter for the Δ*laeA* strain was 1.32-fold higher and similar compared with that of the control strain in the M medium and M medium with 0.6 M potassium chloride, respectively. Complementation of *laeA* rescued the deficient conidia formation of the Δ*laeA* strain, although the Δ*laeA+laeA* strain exhibited 1.26-fold higher conidia formation in the M medium containing 0.6 M potassium chloride.

These results indicate that LaeA is required for asexual development, particularly in the presence of sodium chloride-specific stress versus high osmotic stress. The deficiency in conidia formation was consistent with previous reports that the *laeA* disruption caused a significant defect in conidia formation in *A. oryzae* (18) and *Penicillium chrysogenum* (19). Although LaeA plays a significant role in the sexual development of *A. nidulans* (20), sexual development has not been observed in *A. kawachii*.

### Gene expression related to citric acid production

To identify LaeA-regulated genes related to citric acid production, the expression profiles of *A. kawachii* control and Δ*laeA* strains during citric acid production were compared. These strains were precultured in M medium at 30°C for 36 h and then transferred to CAP medium and further cultured at 30°C for 12 h, at which time the *A. kawachii* control strain vigorously produced citric acid. Next, gene expression profiles were compared using CAGE.

Gene expression change of a total of 9,647 genes was evaluated by CAGE (Data Set S1). The change in gene expression was considered to be statistically significant if the false discovery rate was <0.05 and the log2-fold change was <−0.5 or > 0.5. Using these criteria, a total of 1,248 differentially expressed genes were identified, including 590 upregulated and 658 downregulated genes. Gene ontology (GO) term enrichment analysis of these gene data sets revealed the enrichment of GO terms related to transport and metabolic processes (data not shown); however, it was difficult to interpret the results to explain the reduced citric acid productivity of Δ*laeA* strain.

Next, we mapped the differentially expressed genes, 15 of which were mapped to metabolic pathways related to citric acid production (Fig. 5; Table 1). For example, genes related to the glycerol synthesis [#1 (AKAW_07170) and #12 (AKAW_08691)] and pentose phosphate [#11 (AKAW_00489)] pathways were upregulated by *laeA* disruption, whereas the Embden–Meyerhof–Parnas (EMP) pathway [#2 (AKAW_04737) and #3 (AKAW_03026)] and citrate synthase in TCA or glyoxylate cycles [#4 (AKAW_00170) and #5 (AKAW_06223)] were downregulated by *laeA* disruption. The citrate synthase-encoding gene (#5 [AKAW_06223]) showed most significant reduction in gene expression (0.00038-fold); however, it was not the most highly expressed citrate synthase gene, indicating that the reduced expression level of AKAW_06223 might not explain the deficient citrate productivity.

**Figure 5.**
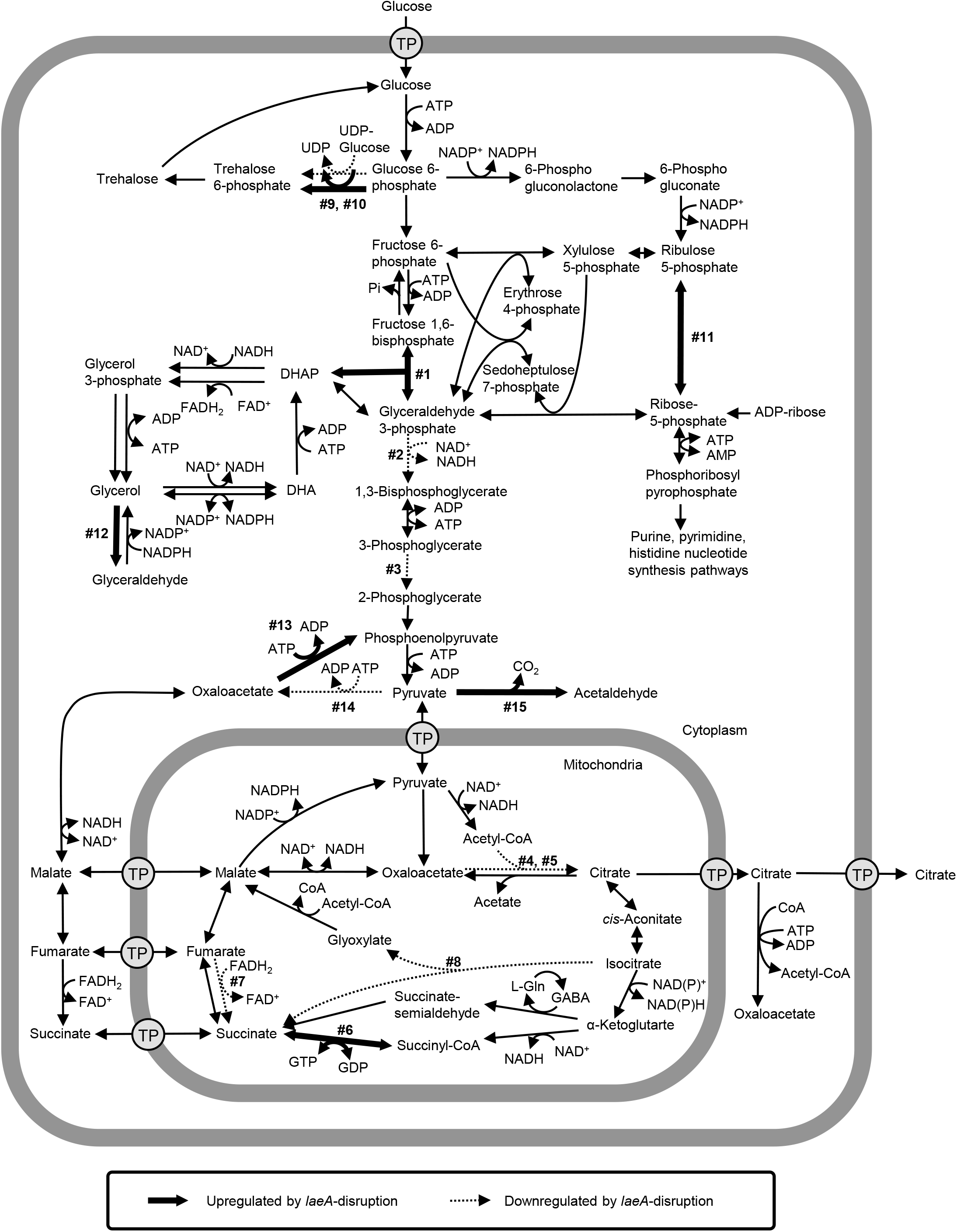
Differentially expressed genes mapped on the proposed metabolic pathways of *A. kawachii*, pathway was constructed based on the metabolic model of *A. niger* and *A. nidulans* (37–39). The glyoxylate shunt was placed in the mitochondria. Bold and dashed arrows indicate downregulated and upregulated reactions due to *laeA* disruption, respectively. Numbers (1~15) next to the arrows correspond to the numbering of locus tags in Table 2. Abbreviations: CoA, coenzyme A; DHA, dihydroxyacetone; DHAP, dihydroxyacetone phosphate; FAD, flavin adenine dinucleotide; Gln, glutamine; TP, trans porter; UDP, uridine diphosphate.

**Table 1.** Citric acid production-related genes among up- and downregulated genes following *laeA* disruption.

| Reaction no.<br>in Fig.5 | Locus tag | Putative function | Fold<br>change | FDR |
| --- | --- | --- | --- | --- |
| #1 | AKAW_07170 | Fructose-bisphosphate aldolase | 16 | 0.0011 |
| #2 | AKAW_04737 | Glyceraldehyde 3-phosphate<br>dehydrogenase | 0.48 | 0.0005 |
| #3 | AKAW_03026 | Phosphoglycerate mutase | 0.58 | 0.0335 |
| #4 | AKAW_00170 | Citrate synthase | 0.53 | 0.0014 |
| #5 | AKAW_06223 | Citrate synthase | 0.00038 | 1.10E-98 |
| #6 | AKAW_07594 | Succinate-CoA ligase (GDP-forming)<br>alpha chain | 2.4 | 0.0052 |
| #7 | AKAW_07352 | Succinate dehydrogenase | 0.62 | 0.0352 |
| #8 | AKAW_00119 | Isocitrate lyase | 0.38 | 1.063E-0<br>5 |
| #9 | AKAW_05189 | Alpha,alpha-trehalosephosphate synthase<br>(UDP-forming) 1 (tpsA, tpsB) | 0.59 | 0.0108 |
| #10 | AKAW_03597 | Alpha,alpha-trehalosephosphate synthase<br>(UDP-forming) 1 (tpsA, tpsB) | 1.8 | 0.0024 |
| #11 | AKAW_00489 | Ribose-5-phosphate isomerase (B) | 2.1 | 0.0035 |
| #12 | AKAW_08691 | Alcohol oxidase | 3.1 | 3.18E-09 |
| #13 | AKAW_02095 | Phosphoenolpyruvate carboxykinase<br>(ATP) | 1.7 | 0.0139 |
| #14 | AKAW_00804 | Pyruvate carboxylase (cytosol) | 0.45 | 0.0168 |
| #15 | AKAW_06947 | Pyruvate decarboxylase | 1.8 | 0.0205 |

Next, we focused on the citric acid export process as it is considered to play a significant role in the high citric acid production of *A. niger* (21–23) (Table 2). Our group previously reported that the two mitochondrial citrate transporters *ctpA* (AKAW_03754) and *yhmA* (AKAW_06280) (24) are involved in the transport of citric acid from mitochondria to cytosol: however, their gene expression levels did not significantly change. Conversely, a significant change in gene expression (0.01-fold reduction) of the putative citrate exporter encoding *cexA* (AKAW_07989) was identified as a citrate exporter from cytosol to extracellular in *A. niger* (15, 16). AKAW_07989 showed the best reciprocal BLAST hit to *A. niger* CexA with 96% identity over 502 amino acid residues, indicating that AKAW_07989 represents CexA in *A. kawachii*.

**Table 2.**
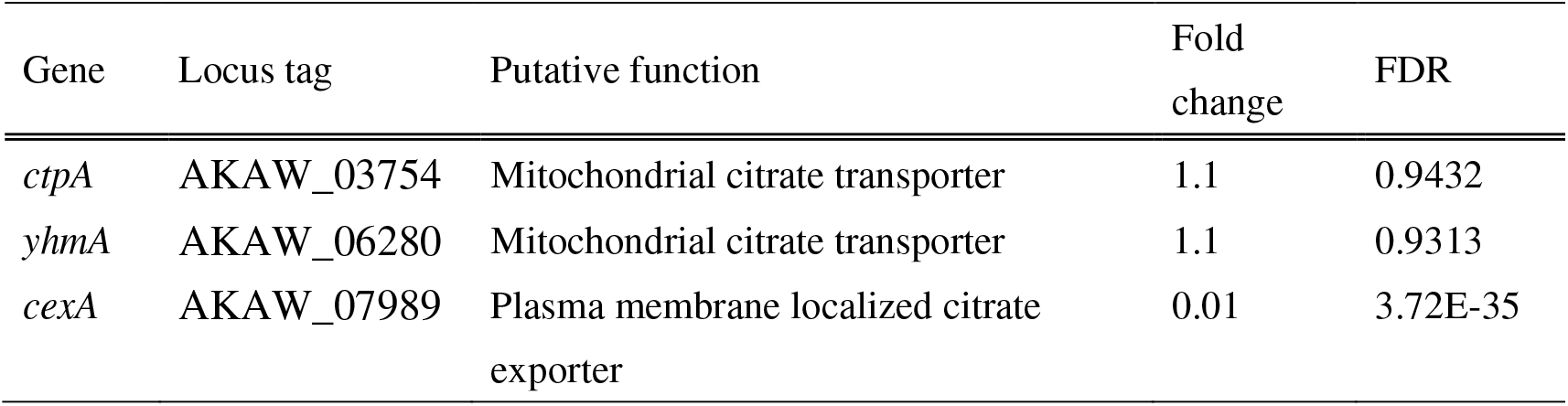
Gene expression change of citric acid excretion-related genes following *laeA* disruption.

Citric acid was the most significantly decreased organic acid in the extracellular fraction resulting from the disruption of *laeA* (Fig. 3A); however, citric acid, malic acid, and oxoglutaric acid productions were reduced at similar levels within the intracellular fraction of the Δ*laeA* strain (Fig. 3B). Thus, we focused on the significantly reduced expression level of *cexA*, which could explain the reduced citric acid accumulation in the extracellular fraction (Table 2).

### Complementation test of *laeA* disruptant by *cexA* overexpression

To examine whether the downregulation of *cexA* was crucial for the decrease in citric acid production caused by *laeA* disruption, we compared the citric acid productivity of control and Δ*laeA* strains with those of Δ*cexA*, Δ*cexA*OE*cexA* (*cexA* overexpression with deletion of *cexA*), and Δ*laeA*OE*cexA* (*cexA* overexpression with deletion of *laeA*) (Fig. 6A). The *gpdA* gene encodes glyceraldehyde-3-phosphate dehydrogenase (25, 26) and was used for overexpression of *cexA* because there was no significant change in *gpdA* gene expression between control and Δ*laeA* strains in CAGE data set (Data Set S1), indicating that the *gpdA* promoter is not controlled by LaeA.

**Figure 6.**
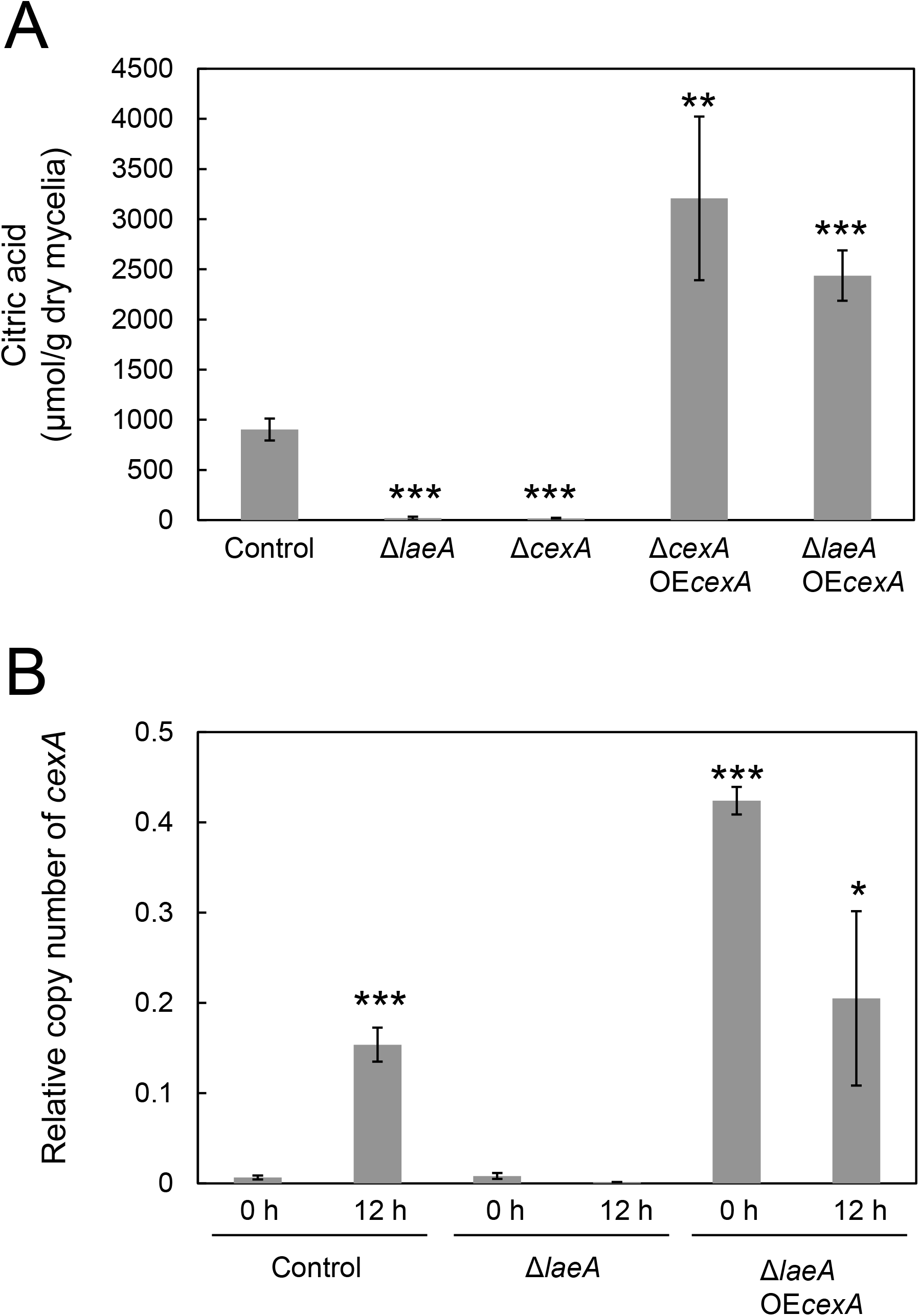
(A) Extracellular citric acid production by *A. kawachii* strains. The control, Δ*laeA*, Δ*cexA*, Δ*cexA*OE*cexA*, and Δ*laeA*OE*cexA* strains were cultured as described. Mean and standard deviation were determined from the results of 3 independent cultivations. Asterisks indicate a statistically significant difference (**, *P* value <0.01; ***, *P* value <0.001; Student’s t-test) from the results for the control strain. (B) Transcriptional levels of *cexA*; control, Δ*laeA*, and Δ*laeA*OE*cexA* strains were precultured in M medium for 36 h, then transferred to CAP medium and further cultured for 0 or 12 h. Mean and standard deviation were determined from the results of 3 independent cultivations. Asterisks indicate a statistically significant difference (*, *P* value < 0.05; ***, *P* value <0.001; Student’s t-test) from the results for the control strain cultured for 0 h.

*A. kawachii* strains were precultured with M medium at 30°C for 36 h, and were then transferred to CAP medium and further cultured at 30°C for 48 h. The culture supernatant was collected as the extracellular fraction. The Δ*cexA* strain exhibited 0.019-fold lower production of extracellular citric acid than the control strain, similar to that observed in the Δ*laeA* strain, whereas the Δ*cexA*OE*cexA* and Δ*laeA*OE*cexA* strains exhibited 3.6-fold and 2.7-fold higher extracellular citric acid production, respectively, than the control strain. There is no statistical difference in citric acid productivity between strains Δ*cexA*OE*cexA* and Δ*laeA*OE*cexA*, indicating that the overexpression of *cexA* alone was sufficient to rescue the citric acid productivity deficit in the Δ*laeA* strain.

### Transcriptional levels of *cexA* in control, Δ*laeA*, and Δ*laeA*OE*cexA*

Because Δ*laeA*OE*cexA* exhibited 2.7-fold higher citric acid productivity than the control strain (Fig. 6A), we determined gene expression levels of *cexA* in control, Δ*laeA*, and Δ*laeA*OE*cexA*. Strains were precultured in M medium at 30°C for 36 h, and then transferred to CAP medium. The time point at 36 h of precultivation just before transfer was defined as 0 h (i.e., starting time). We compared the gene expression level of *cexA* at 0 and 12 h of cultivation in CAP medium via quantitative RT-PCR analysis (Fig. 6B). The control strain showed a 24-fold greater expression level of *cexA* at 12 h than that at 0 h, whereas the Δ*laeA* strain showed similar expression levels of *cexA* at 0 and 12 h. These results indicate that *cexA* expression is induced under condition of citric acid production via LaeA.

Control and Δ*laeA*OE*cexA* strains showed similar expression levels of *cexA* at 12 h; however, Δ*laeA*OE*cexA* exhibited a 66-fold higher expression level of *cexA* than the control strain at 0 h, likely because the *gpdA* promoter is active in M as well as CAP media. Thus, the higher citric acid productivity of Δ*laeA*OE*cexA* (Fig. 6A) could be owing to a higher gene expression level of *cexA* when beginning culture in CAP medium.

### Histone trimethylation level in the *cexA* promoter region

LaeA is believed to widely regulate gene expression via controlling methylation level of histones (11–13). To determine the mechanism underlying LaeA-dependent *cexA* expression through histone methylation, we performed ChIP-qPCR analysis of histone H3, histone histone H3 trimethyl K4 (H3K4 me3), and histone H3 trimethyl K9 (H3K9 me3) in control and Δ*laeA* strains. H3K4 me3 and H3K9 me3 are known to be euchromatin and heterochromatin markers, respectively.

The histone H3 occupancy at the *cexA* promoter did not change between control and Δ*laeA* strains (Fig. 7A). Alternatively, the euchromatin marker H3K4 me3 at the *cexA* promoter was decreased to a level similar to that of the negative control (i.e., normal anti-mouse IgG) in Δ*laeA* (Fig. 7B). In addition, the heterochromatin marker H3K9 me3 was greatly enriched in Δ*laeA* compared with that in the control strain (Fig. 7C). These results indicated that LaeA controls *cexA* expression via modulation of euchromatin/heterochromatin ratios at the *cexA* promoter region in *A. kawachii*.

**Figure 7.**
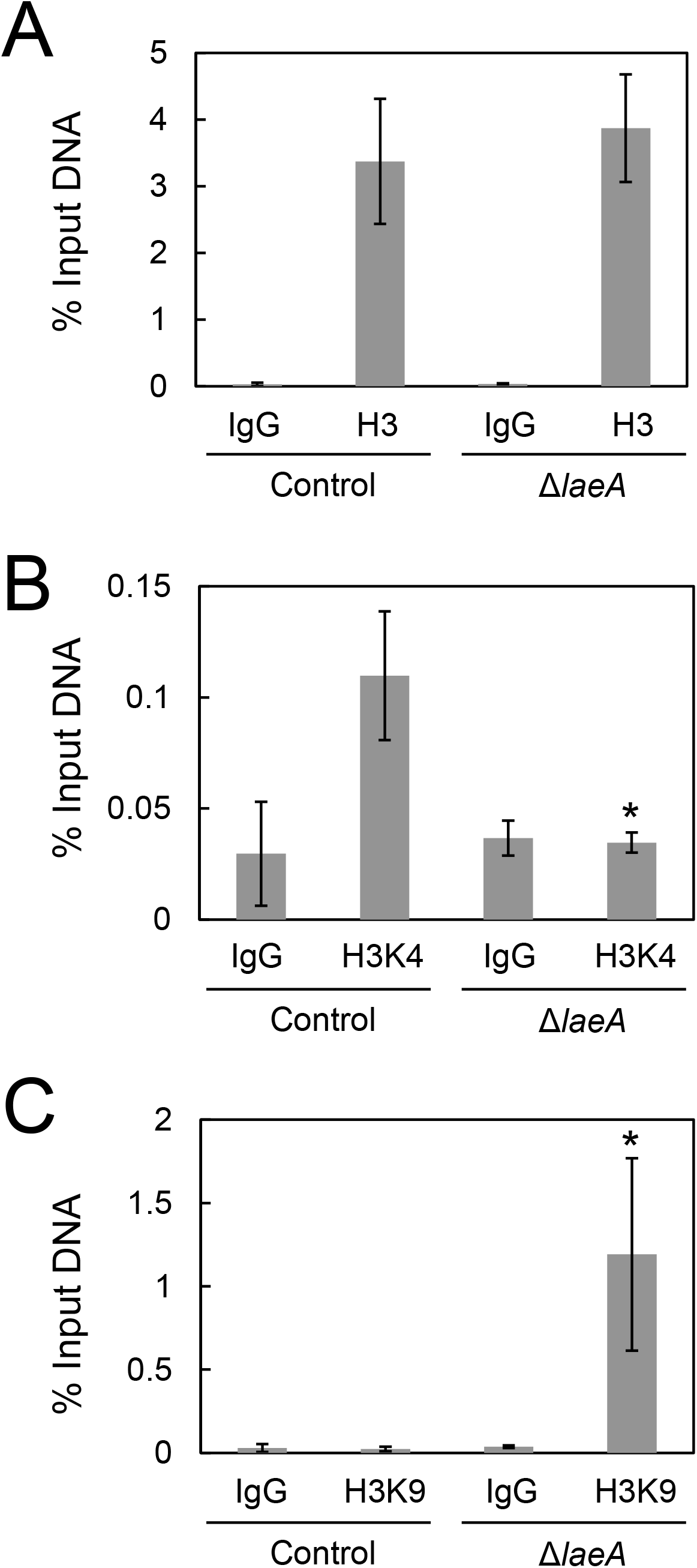
ChIP-qPCR of *cexA* promoter region; control and Δ*laeA* were cultured as described. (A) Histone H3, (B) histone H3K4 me3, and (C) histone H3K9 me3 occupancies of the *cexA* promoter region were investigated. The mean and standard deviation were determined from the results of 3 independent cultivations. Asterisks indicate a statistically significant difference (*, *P* value < 0.05; Student’s t-test) from the results for the control strain.

Although details concerning the molecular function of LaeA remain unclear (11–13), LaeA is known to counteract the heterochromatinization of the promoter region of secondary metabolite gene clusters by histone H3K9 methylation via heterochromatin protein 1 (HepA) and a H3K9 methyltransferase (ClrD) in *A. nidulans* (27). Our finding indicated that the euchromatin structure of the *cexA* promoter nearly disappeared with *laeA* disruption, and therefore the heterochromatin level of the *cexA* promoter might be enriched by HepA (AKAW_02119) and ClrD (AKAW_07568) orthologs in *A. kawachii*. Gene expression levels of AKAW_02119 and AKAW_07568 were not significantly altered in Δ*laeA* (Data Set S1), but loss of LaeA might affect histone modification balance and euchromatin/heterochromatin ratios. The molecular mechanism of LaeA-dependent histone modification should be confirmed through additional experiments. In addition, whether gene expression of *cexA* requires a specific DNA-binding transcriptional factor remains unclear and should be further studied.

In conclusion, LaeA plays a significant role in citric acid production in *A. kawachii* by controlling *cexA* expression via histone modification at the *cexA* promoter region. Because *A. kawachii* is widely used in the production of shochu and elsewhere in the fermentation industry, our findings are expected to enhance the understanding of citric acid production mechanism(s) and facilitate optimization of strategies for controlling *A. kawachii* activity.

## MATERIALS AND METHODS

### Strains and growth conditions

The *Aspergillus kawachii* strains used in this study are listed in Table S1, and strain SO2 (28) was used as the parental strain.

For construction and characterization, the strains were grown in minimal medium (1% [wt/vol] glucose, 0.6% [w/v] NaNO_3_, 0.052% [w/v] KCl, 0.052% [w/v] MgSO_4_⋅7H_2_O, 0.152% [w/v] KH_2_PO_4_, plus Hutner’s trace elements [pH 6.5]). Medium was adjusted to the required pH using HCl or NaOH. For the cultivation of *sC*^−^ and *argB*^−^ strains, 0.02% (w/v) methionine and/or 0.211% (w/v) arginine were added to M medium, respectively.

To evaluate acidification occurring on agar medium, strains were grown in YPD with methyl red (2% [w/v] glucose, 1% [w/v] yeast extract, 2% [w/v] peptone, and 2% methyl red solution) prepared as follows: 100 mg methyl red (Nakalai Tesque, Kyoto, Japan) was dissolved in 100 ml ethanol and titrated by 0.1% (w/v) NaOH solution until observation of an obvious color change from red to yellow; the solution was then sterile filtered (0.2 μm pore size; Toyo Roshi Kaisha, Japan).

To investigate citric acid productivity, *A. kawachii* strains were also grown in CAP medium (10% [w/v] glucose, 0.3% [w/v] (NH_4_)_2_SO_4_, 0.001% [w/v] KH_2_PO_4_, 0.05% [w/v] MgSO_4_⋅7H_2_O, 0.000005% [w/v] FeSO_4_⋅7H_2_O, 0.00025% [w/v] ZnSO_4_⋅5H_2_O, 0.00006% [w/v] CuSO_4_⋅5H_2_O [pH 4.0]). CAP medium was adjusted to the required pH with HCl.

### Construction of putative methyltransferase gene-disruptant strain

The *laeA*, *laeA2*, and *laeA3* genes were disrupted in *A. kawachii* SO2 (28) by insertion of the *argB* gene. A gene replacement cassette encompassing the homology arm at the 5’ end of the putative methyltransferase genes, an *argB* selection marker, and homology arm at the 3’ end of the putative methyltransferase genes were constructed by recombinant PCR using the primer pairs AKlaeX-FC/AKlaeX-R1, argB-F2/argB-R2, and AKlaeX-F3/AKlaeX-RC, respectively (where “X” indicates A, A2 or A3) (Table S2). For amplification of the *argB* gene, the pDC1 plasmid was used as template DNA (29). For amplification of other DNA fragment(s), *A. kawachii* IFO 4308 wild type genomic DNA was used as template DNA. The resultant DNA fragment amplified with primers AKlaeX-F1 and AKlaeX-R3 was used to transform *A. kawachii* SO2, yielding strains Δ*sC*Δ*laeA*, Δ*sC*Δ*laeA2*, and Δ*sC*Δ*laeA3*. M agar plates lacking arginine were used for the selection of transformants. Introduction of the *argB* gene into each methyltransferase gene locus was confirmed by PCR using the primer pairs AKlaeX-FC and AKlaeX-RC (Fig. S1).

The SO2 strain was transformed using the *argB* gene cassette to employ the same auxotrophic genetic background strains for comparative study. This *argB* gene cassette was generated with PCR using *A. kawachii* genomic DNA and pDC1 as template DNA and was used to transform the SO2 strain, yielding the Δ*sC* strain (Table S1). Transformants were selected on M agar medium lacking arginine.

### Complementation of the *laeA*-disruptant strain

For complementation analysis of the *laeA* disruptant using wild type *laeA*, a gene replacement cassette encompassing a homology arm at the 5’ end of *laeA*, wild type *laeA*, *sC* selection marker, and homology arm at the *argB* locus was constructed with recombinant PCR using the primer pairs AKlaeA-FC/AKlaeAcomp-R1 and AKlaeAcomp-F2/argB-R2, respectively (Table S2). For amplification of DNA fragments, *A. kawachii* IFO 4308 wild type genomic DNA and a plasmid carrying tandemly connected *sC* and *argB* were used as template DNA. The resultant DNA fragment amplified with primers AKlaeA-F1/argB-R2 was used to transform the *laeA* disruptant, yielding the Δ*laeA*+*laeA* strain. Transformants were selected on M agar medium lacking methionine. Introduction of the *laeA* and *sC* genes into the target locus was confirmed by PCR using primers AKlaeA-FC and argB-R2 (Fig. S2).

The Δ*sC*Δ*laeA* strain was transformed using the *sC* gene cassette for use of the same auxotrophic genetic background strains for the comparative study. The *sC* gene cassette was generated by PCR using *A. kawachii* genomic DNA as the template and primers sC-comp-F and sC-comp-R (Table S2), and was used to transform the Δ*sC*Δ*laeA* strain and yield the Δ*laeA* strain (Table S1). Transformants were selected on M agar medium lacking methionine.

### Construction of putative citrate exporter gene-disruptant strain

*cexA* was disrupted in *A. kawachii* SO2 (28) by insertion of the *argB* gene. A gene replacement cassette encompassing the homology arm at the 5’ end of the *cexA*, *argB* selection marker, and homology arm at the 3’ end of *cexA* was constructed using recombinant PCR with the primer pairs AKcexA-FC/AKcexA-R1, AKcexA-F2/AKcexA-R2, and AKcexA-F3/AKcexA-RC, respectively (Table S2). For amplification of the *argB* gene, the plasmid pDC1 was used as the template DNA (29). For amplification of the other DNA fragment, *A. kawachii* IFO 4308 wild type genomic DNA was used as a template. The resultant DNA fragment was amplified with primers AKcexA-F1 and AKcexA-R3 and was used to transform *A. kawachii* SO2 and yield strain Δ*sC*Δ*cexA*. M agar plates lacking arginine were used for selection of transformants. Introduction of *argB* into the *cexA* locus was confirmed by PCR using the primer pair AKcexA-FC and AKcexA-RC (Fig. S3). After confirmation of gene disruption, the Δ*sC*Δ*cexA* strain was transformed with the *sC* gene cassette to use the same auxotrophic genetic background strains for comparative study. This cassette was synthesized by PCR using primers sC-comp-F and sC-comp-R and *A. kawachii* genomic DNA as template (Table S2). Transformants were selected on M agar medium lacking methionine.

### Construction of the putative citrate exporter-overexpression strain

The plasmid pGS-PgpdA (30) was used for overexpression of *cexA*. The *cexA* was amplified by PCR with primers pGSG-cexA-inf-F/pGSG-cexA-inf-R using *A. kawahcii* genomic DNA as template (Table S2). The amplicon was inserted into the SalI site of pGS-PgpdA, thereby yielding pGS-PgpdA-cexA, which was used to transform strains Δ*sC*Δ*cexA* and Δ*sC*Δ*laeA*, yielding Δ*cexA*OE*cexA* and Δ*laeA*OE*cexA*, respectively (Table S1).

### Measurement of extracellular and intracellular organic acids

Levels of extracellular and intracellular organic acids were measured as described previously (24). Briefly, 2 × 10^7^ conidia cells of *A. kawachii* were inoculated into 100 ml M medium, precultured with shaking (180 rpm) at 30°C for 36 h, and then transferred into 50 ml CAP medium and further cultured with shaking (163 rpm) at 30°C for 48 h. The culture supernatant was harvested as the extracellular fraction. Mycelia were used for preparation of the intracellular fraction using a hot-water extraction method (31) with modifications. The freeze-dried as well as wet mycelial weights were measured as extracellular and intracellular fractions, respectively. Wet mycelia were ground into a powder using mortar and pestle in the presence of liquid nitrogen, and then dissolved in 10 ml of hot water (80°C) per 1 g of mycelial powder, vortexed, and centrifuged at 18,800 × *g* at 4°C for 30 min. The supernatant was taken as the intracellular fraction. Extracellular and intracellular fractions were filtered through a PTFE filter (0.2 μm pore size; Toyo Roshi Kaisha) and analyzed with HPLC on a Prominence HPLC system (Shimadzu, Kyoto, Japan) equipped with a CDD-10AVP conductivity detector (Shimadzu). The organic acids were separated using two tandem Shimadzu Shim-pack SCR-102H columns (internal diameter 300 by 8 mm; Shimadzu) at 50°C using 4 mM *p*-toluenesulfonic acid monohydrate as the mobile phase at a flow rate of 0.8 ml/min. The flow rate of the post-column reaction solution (4 mM *p*-toluenesulfonic acid monohydrate, 16 mM bis-Tris, and 80 μM EDTA) was 0.8 ml/min.

### Cap-analysis gene expression (CAGE) analysis

Total RNA was extracted from mycelia. Briefly, 2 × 10^7^ conidia cells of the *A. kawachii* strains were inoculated into 100 ml of M medium, precultured with shaking (180 rpm) at 30°C for 36 h, and then transferred to 50 ml of CAP medium and further cultured with shaking (163 rpm) at 30°C for 12 h. The mycelia were ground to a powder as described above. Then, RNA was extracted using RNAiso Plus reagent (Takara Bio, Shiga, Japan). RNA samples were treated with the SV total RNA isolation system (Promega, Madison, WI) according to the manufacturer’s protocol.

Library preparation, sequencing, and data analysis for CAGE (32) were performed by K.K. DNAFORM (Kanagawa, Japan). All CAGE-seq experiments were performed three times with RNA samples obtained from independently prepared mycelia. First-strand cDNAs were transcribed to the 5’ end of capped RNAs, attached to CAGE barcode tags and these tags were sequenced using the NextSeq 500 system (Illumina, San Diego, CA) and mapped to the *A. kawachii* IFO 4308 genome (33) using BWA software (v0.5.9) after discarding ribosomal or non-A/C/G/T base containing RNAs. For tag clustering, CAGE-tag 5’ coordinates were input for RECLU clustering (34). The criteria for linking transcriptional start sites and predicted coding sequences were within 600 bp upstream or downstream of the predicted start codon. Triplicate data were analyzed and the expression ratio was also calculated as the log (base 2) ratio through the RECLU pipeline.

CAGE data were deposited in the Gene Expression Omnibus under accession number GSE135849.

### Transcriptional analysis

For RNA extraction from mycelia, conidia (2 × 10^7^ cells) of the *A. kawachii* control, Δ*laeA*, and Δ*laeA*OE*cexA* strains were inoculated into 100 ml M medium and cultured with shaking (180 rpm) for 36 h at 30°C. After incubation, mycelia were collected and divided into two half portions and transferred individually to CAP medium and cultured with shaking (163 rpm) for 12 h at 30°C. Mycelia were ground as described above and RNA was extracted using RNAiso Plus (Takara Bio), then cDNA was synthesized from total RNA using a PrimeScript Perfect Real-Time Reagent Kit (Takara Bio) according to manufacturer’s protocols. Real-time RT-PCR was performed using a Thermal Cycler Dice Real-Time System MRQ (Takara Bio) with SYBR Premix Ex Taq II (Tli RNaseH Plus) (Takara Bio). The following primer sets were used: AKcexA-RT-F and AKcexA-RT-R for *cexA* and AKactA-RT-F and AKactA-RT-R for *actA* (Table S2).

### ChIP-qPCR

*A. kawachii* conidia were cultured as described above. ChIP was performed as previously described (35) using normal anti-mouse IgG as a negative control (Cosmo Bio, Tokyo, Japan), as well as anti-histone H3 (Medical and Biological Laboratories, Nagoya, Japan), anti-H3K4 me3 (Medical and Biological Laboratories), and anti-H3K9 me3 antibodies (Medical and Biological Laboratories). Two µg antibodies were used with 200 mg of total protein in each ChIP experiment. DNA quantification was measured with real-time qPCR using SYBR Premix Ex Taq II (Tli RNaseH Plus) (Takara Bio) and the primer set cexA-ChIP-F and cexA-ChIP-R (Table S2). Positions of the primers relative to the ATG site of *cexA* were +2 to +26 for cexA-ChIP-F and +238 to +257 for cexA-ChIP-R. Relative amounts of DNA (i.e. percent input DNA) were calculated by dividing immunoprecipitated DNA by input DNA.

## ACKNOWLEDGMENTS

We thank Kenta Hamada and Aoi Miyamoto for helpful discussion and technical supports. This work was supported by JSPS KAKENHI grants (18K05394 and 19K05773) and a grant from Noda Institute for Scientific Research. C.K. was supported by a Grant-in-Aid for JSPS Research Fellows (17J02753).

